# Counting rare *Wolbachia* endosymbionts using digital droplet PCR

**DOI:** 10.1101/2024.12.10.627731

**Authors:** Alphaxand K. Njogu, Francesca Logozzo, William R. Conner, J. Dylan Shropshire

## Abstract

*Wolbachia* is the most widespread animal-associated intracellular microbe, living within the cells of over half of insect species. Since they can suppress pathogen replication and spread rapidly through insect populations, *Wolbachia* is at the vanguard of public health initiatives to control mosquito-borne diseases. *Wolbachia*’s abilities to block pathogens and spread quickly are closely linked to their abundance in host tissues. The most common method for counting *Wolbachia* is quantitative polymerase chain reaction (qPCR), yet qPCR can be insufficient to count rare *Wolbachia*, necessitating tissue pooling and consequently compromising individual-level resolution of *Wolbachia* dynamics. Digital droplet PCR (ddPCR) offers superior sensitivity, enabling the detection of rare targets and eliminating the need for sample pooling. Here, we report three ddPCR assays to measure total *Wolbachia* abundance, *Wolbachia* abundance adjusted for DNA extraction efficiency, and *Wolbachia* density relative to host genome copies. Using *Drosophila melanogaster* with *w*Mel *Wolbachia* as a model, we show these ddPCR assays can reliably detect as few as 7 to 12 *Wolbachia* gene copies in a 20 μL reaction. The designed oligos are homologous to sequences from at least 106 *Wolbachia* strains across Supergroup A and 53 host species from the *Drosophila*, *Scaptomyza*, and *Zaprionus* genera, suggesting broad utility. These highly sensitive ddPCR assays are expected to significantly advance *Wolbachia*-host interactions research by enabling the collection of molecular data from individual insect tissues. Their ability to detect rare *Wolbachia* will be especially valuable in applied and natural field settings where pooling samples could obscure important variation.

## Importance

Mosquitoes pose a significant threat to human health, but *Wolbachia* bacteria have emerged as a promising tool to combat the transmission of the deadly diseases they vector. These bacteria can spread swiftly through insect populations and confer resistance to mosquito-borne diseases including dengue and Zika viruses, effectively blocking their transmission to humans. *Wolbachia* abundance within an insect at least partly explains the strength of the traits that make *Wolbachia* valuable for disease control, making precise measurement essential for public health applications. However, traditional molecular assays can lack the sensitivity needed for accurate, individual-level quantification for rare *Wolbachia*. Here, we present three highly sensitive digital droplet PCR (ddPCR) assays for *Wolbachia* detection, offering superior sensitivity compared to existing methods. These assays will be useful for studies that measure *Wolbachia* abundance and related phenotypes in individual insects, providing enhanced resolution and improving efforts to characterize the mechanisms that govern phenotypic variation.

## Introduction

Endosymbiotic relationships, where one organism resides within the cells of another, are some of the most intimate biological partnerships. The alphaproteobacteria *Wolbachia* is among the most widespread endosymbionts, living in the cells of over half of all insect species, numerous other arthropods, and filarial nematodes (Taylor et al., 2005; Weinert et al., 2015). *Wolbachia*’s success can be explained by efficient maternal transmission, effects on host reproduction that favor females with *Wolbachia*, and fitness advantages conferred by *Wolbachia* to its host (Hague et al., 2022; Hoffmann et al., 1990). Reproductive effects include killing or feminizing males (Hurst et al., 1999; Rigaud et al., 1991), inducing thelytokous parthenogenesis (Ma and Schwander, 2017), and causing a conditional male sterility called cytoplasmic incompatibility (Yen and Barr, 1973). Fitness benefits include nutritional supplementation (Hickin et al., 2022; Newton and Rice, 2020) and protection from pathogens like viruses (Hedges et al., 2008; Teixeira et al., 2008), fungi (Perlmutter et al., 2023), and protists (Moreira et al., 2009). In addition to ensuring *Wolbachia*’s prevalence in natural populations, these traits make *Wolbachia* a valuable tool to control diseases transmitted by insects. For instance, cytoplasmic incompatibility helps *Wolbachia* spread through *Aedes aegypti* mosquito populations (Hoffmann et al., 2011) where the bacteria can inhibit viral replication and reduce the transmission of dengue and Zika viruses to humans (Hoffmann et al., 2024; Lenharo, 2023; Simmons et al., 2024; Utarini et al., 2021).

*Wolbachia*-associated traits, while crucial for their widespread distribution, exhibit significant variation in strength. A common hypothesis suggests that *Wolbachia*-induced traits are stronger when there are more *Wolbachia* cells within each host cell. This is supported by numerous studies demonstrating a link between *Wolbachia* abundance and the strength of cytoplasmic incompatibility and pathogen inhibition (Chrostek et al., 2013; Clancy and Hoffmann, 1998; Clark et al., 2003; Martinez et al., 2015, 2014; Osborne et al., 2012, 2009; Shropshire et al., 2022; Veneti et al., 2003). Despite some exceptions to these trends (reviewed in Shropshire et al., 2020), *Wolbachia* abundance measures remain valuable for their simplicity and application when the genetic basis of a trait is unknown. Quantitative polymerase chain reaction (qPCR) is a widely used method for measuring *Wolbachia* abundance. By tracking fluorescence accumulation during PCR cycles, qPCR enables the quantification of target DNA sequences. This can be achieved through absolute *Wolbachia* quantification (titer), which involves comparing results to a standard curve, or relative quantification, which normalizes *Wolbachia* abundance against a host gene (density). Notably, relative quantification relies on assumptions about consistent host genome copy number across treatment groups, which may not always hold true (Christensen et al., 2019). Therefore, careful consideration should be given when choosing between titer and density assays.

While qPCR-based assays are widely used to quantify *Wolbachia* abundance, they can suffer from accuracy and sensitivity limitations. Suboptimal PCR amplification efficiency and variable background noise can lead to underestimation of target abundance and limit the sensitivity of an assay (Forootan et al., 2017). Although qPCR efficiency can and should be optimized (Bustin et al., 2009), sample-specific biochemical variations can still lead to inconsistent amplification and inaccurate results. Digital droplet PCR (ddPCR) offers a more robust and sensitive alternative (Baker, 2012). In ddPCR, a PCR reaction is partitioned into thousands of nanoliter-sized droplets, PCR is performed, and droplets are then individually passed through a microfluidics fluorescence detector to count droplets with and without the target. Applying Poisson statistics to the droplet data allows for absolute quantification of target molecules without the need for a standard curve. As an end-point PCR assay, ddPCR is robust to variation in PCR efficiency compared to qPCR where quantification assumes the target doubles every PCR cycle. Moreover, since ddPCR can detect targets in droplets containing only a single molecule, its limit of detection (LoD) is primarily determined by the number of positive droplets detected in the reaction (as low as 0.001%) and the rate of false-positive droplets in negative controls.

To date, ddPCR has been used to quantify *w*Cle, *w*Pip, and *w*AlbB *Wolbachia* strains in *Cimex lectularius* bed bugs and *Ae. albopictus* mosquitoes (Fisher et al., 2019; Hickin et al., 2022; Kakumanu et al., 2024; Kilpatrick et al., 2024). To expand the applicability of ddPCR to a broader range of *Wolbachia* strains and host systems, we developed singleplex and duplex ddPCR assays targeting: *ftsZ*, a single-copy gene of *Wolbachia*; *mid1*, an ultraconserved element of the *Drosophila* genus; and a commercially available DNA spike-in control. The singleplex *ftsZ* assay, designed to be compatible with 106 *Wolbachia* strains from supergroup A, allows for accurate titer calculations. The duplex *ftsZ*/*mid1* and *ftsZ*/spike assays enable the determination of *Wolbachia* density and absolute titer, respectively. Using the *w*Mel *Wolbachia* of *D. melanogaster* as a model, we demonstrate a LoD of 7 to 12 *Wolbachia* gene copies per reaction, and similar LoDs for *mid1* and spike-in control copies. These assays are well-suited for analyzing complex samples, low biomass samples, and samples with rare *Wolbachia*.

## Results

### *Wolbachia* absolute abundance from singleplex ddPCR

The *Wolbachia* gene *ftsZ* is a common target for qPCR-based estimates of *Wolbachia* abundance since it is a conserved, single-copy gene across *Wolbachia* genomes (e.g. Shropshire et al., 2022). To design primers and probes for *ftsZ*, we first obtained *ftsZ*-annotated sequences from a large collection of *Wolbachia* genomes, generated consensus sequences from a multiple sequence alignment (MSA), and masked variable nucleotides. Using Primer3Web and NCBI Primer Blast, we designed a forward primer, reverse primer, and FAM-labeled probe (see details in Materials & Methods). To refine our oligo selection, we iteratively narrowed the list by removing sequences distantly related to *w*Mel *Wolbachia* in *D. melanogaster*, ultimately arriving at suitable oligos based on 106 *ftsZ* sequences from 106 supergroup A *Wolbachia* strains (**Fig 1**; **Data S1**).

**Figure 1.**
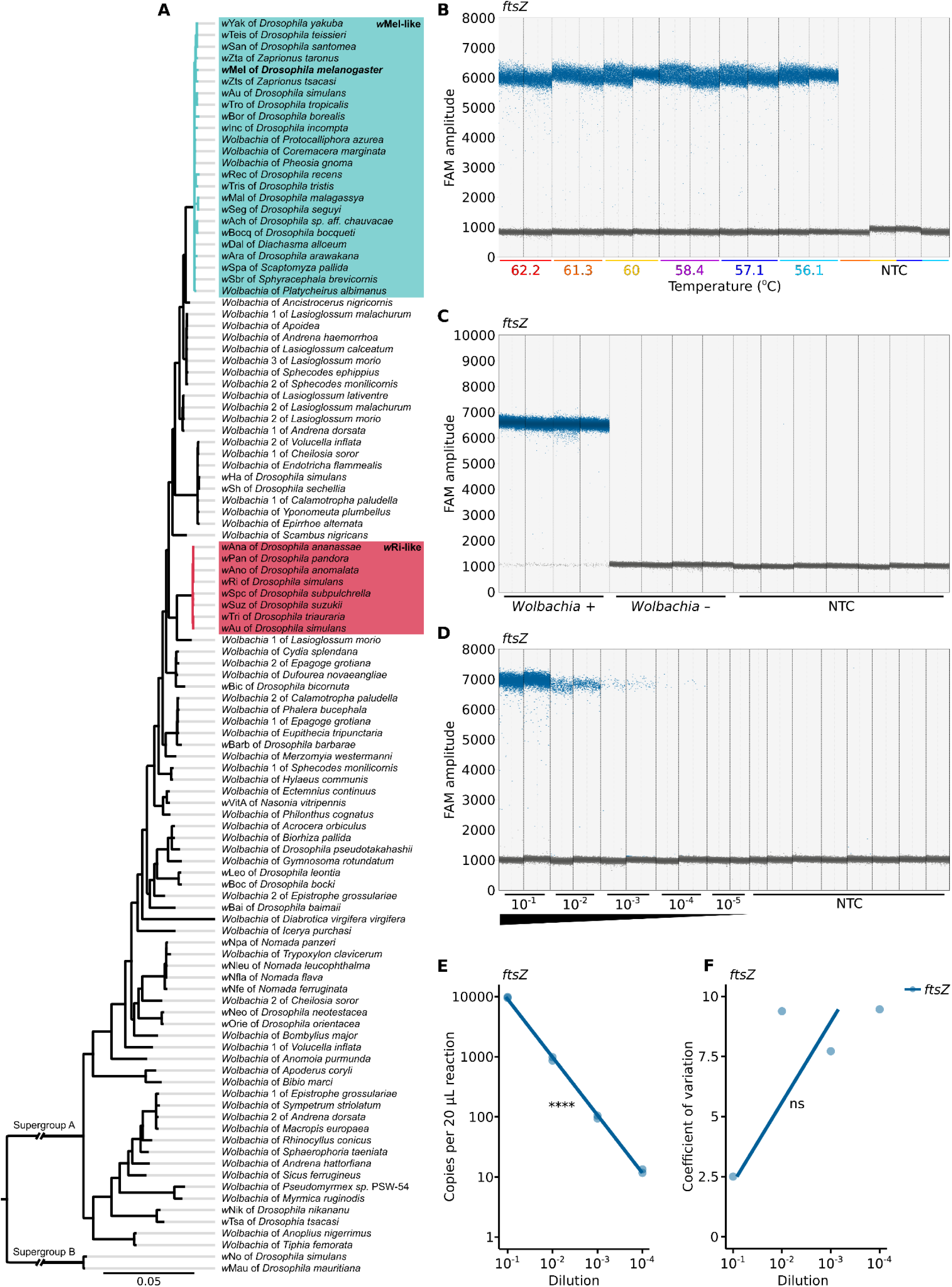
A specific and sensitive *ftsZ* ddPCR assay for counting supergroup A *Wolbachia*. **(A)** Phylogenetic tree of 106 supergroup A *Wolbachia* strains, including outgroups (supergroup B). The *ftsZ* ddPCR primers and probe were designed based on a multiple sequence alignment of these strains (**Data S1**; **Table S2**). Key strains are highlighted: *w*Mel-like (teal) (Shropshire et al., 2024) and *w*Ri-like (light red) (Turelli et al., 2018). The tree was constructed using 69 single-copy genes (50,421 nucleotides) with a posterior probability threshold of 0.95. Branch lengths are substitutions per site. **(B)** *ftsZ* detection is consistent across a range of annealing temperatures (62.2°C to 56.1°C). **(C)** *ftsZ* detection is specific to *Wolbachia*-positive *D. melanogaster* samples. **(D)** The number of *ftsZ*-positive droplets declines across a dilution series. **(B-D)** Each dot represents a droplet, with gray and blue indicating negative and positive, respectively. Samples are separated by vertical dotted lines along the x-axis. **(E)** *ftsZ* concentration measured by ddPCR strongly correlates with the dilution series. **(F)** Variation between technical replicates is not significantly correlated with the dilution series. Statistical significance: *P* > 0.05 (ns), *P* ≤ 0.05 (*), *P* ≤ 0.01 (**), *P* ≤ 0.001 (***), *P* ≤ 0.0001 (****). All statistical tests are Pearson’s product-moment correlations. The raw data is available in **Data S3**.

We used the *w*Mel *Wolbachia* strain of *D. melanogaster* to evaluate *ftsZ*-ddPCR assays. This system is the canonical model for investigating *Wolbachia*-host interactions (Wu et al., 2004), and the *w*Mel strain is widely used in initiatives to control mosquito-borne diseases (e.g. Utarini et al., 2021). We evaluated *ftsZ* oligo performance across a 6°C temperature range (56.2°C to 62.2°C), expecting the optimal annealing temperature to correspond to high fluorescence amplitude on the FAM channel. We observed only minor variation across temperatures, with the lowest FAM amplitude at 62.2°C (*x̄* = 5959) and the highest at 60°C (*x̄* = 6082; **Fig 1B**). Fluorescence amplitudes of negative droplets that do not contain the target were consistent across groups (*x̄_min_* = 827; *x̄_max_*= 838; **Fig 1B**). Therefore, we considered all annealing temperatures to be suitable for differentiating positive and negative droplets, and we selected 60^°C^ for subsequent analyses. To determine if *ftsZ*-ddPCR assays are specific to *Wolbachia*, we subjected DNA from *w*Mel-bearing *D. melanogaster*, *Wolbachia*-free *D. melanogaster*, and no template controls (NTCs) to *ftsZ*-ddPCR assays. Positive droplets were detected in DNA from *w*Mel-bearing flies and no more than 1 in 17,795 positive droplets from *Wolbachia*-free flies or NTCs, confirming that the *ftsZ* ddPCR assay is specific to *Wolbachia* (**Fig 1C**).

To evaluate the sensitivity and precision of the *ftsZ*-ddPCR assay, we measured *ftsZ* abundance across a 1:10 dilution series of DNA from *w*Mel-bearing *D. melanogaster*. We detected *ftsZ* across four dilution factors, from 10^−1^ to 10^−4^ (95% confidence interval (CI) [473, 498] to [0.22, 1.01] *ftsZ* copies/μL, respectively; **Fig 1D**). A strong correlation between *ftsZ* concentration and dilution factor indicates sensitive and accurate quantification (95% CI [0.992, 1] Pearson’s *r^2^*, *P* = 7.1e-10; **Fig 1E**). Moreover, while variation between technical replicates is higher at lower dilutions, this relationship is not significant (95% CI [0.54, 0.99] Pearson’s *r^2^*, *P* = 0.23; **Fig 1F**), suggesting high precision across concentrations. To determine the LoD, we compared these results to those from eight NTCs. We detected four false-positive droplets among 132,579 total droplets, with three of these occurring in a single reaction with a low estimated *ftsZ* concentration (95% CI [0.05, 0.60] *ftsZ* copies/μL). Given that we detected *ftsZ* from *w*Mel DNA down to this concentration range, we estimate the LoD of the *ftsZ*-ddPCR assay to be approximately 12 *ftsZ* copies per 20 μL reaction.

### Measuring *Wolbachia* density with ddPCR

*Wolbachia* density is defined as the ratio of *Wolbachia* genomes to host genomes. To accurately quantify both, we paired the *ftsZ*-ddPCR assay targeting *Wolbachia* with a new *mid1*-ddPCR assay targeting *mid1*, a single-copy “ultraconserved element” in the *Drosophila* genus (Kern et al., 2015). We designed primers and a HEX-labeled probe for *mid1* based on a multiple sequence alignment of 50 *Drosophila*, 1 *Scaptomyza*, and 2 *Zaprionus* species (**Table S3**; **Data S2**). The singleplex *mid1*-ddPCR assay HEX amplitude was lowest at 62.2°C (*x̄* = 13533), highest at 58.4°C (*x̄* = 13854), and negative droplet amplitude was consistent across temperatures (*x̄_min_* = 1121; *x̄_max_* = 1162; **Fig 2A**). We selected 60°C for subsequent reactions, aligning it with the annealing temperature chosen for *ftsZ*. We successfully detected *mid1* across four dilution factors, from 10^−1^ to 10^−4^ (95% CI [272, 291] to [0.17, 0.94] *mid1* copies/μL, respectively; **Fig 2B**). A strong correlation between *mid1* concentration and dilution factor demonstrates accurate and sensitive quantification (95% CI [0.878, 0.996] Pearson’s *r^2^*, *P* = 7.1e-10), and low technical variation between replicates across concentrations demonstrates high precision (95% CI [0.53, 0.99] Pearson’s *r^2^*, *P* = 0.2).

**Figure 2.**
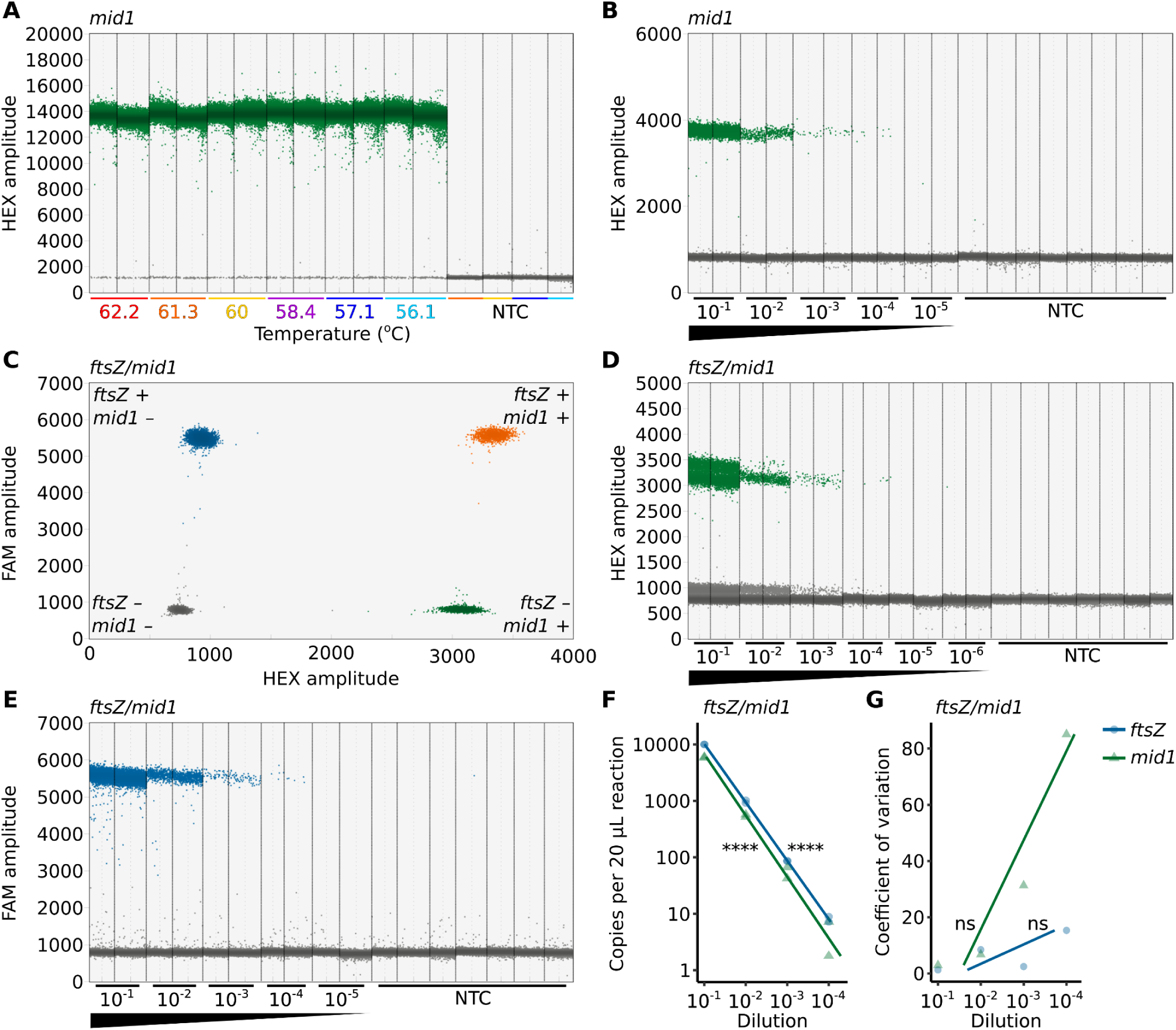
A sensitive *ftsZ*/*mid1* ddPCR assay for measuring *Wolbachia* density in host cells. **(A)** *mid1* detection is consistent across a range of annealing temperatures (62.2°C to 56.1°C). **(B)** The number of *mid1*-positive droplets declines across a dilution series in a mid1 ddPCR assay. **(C)** In duplex *ftsZ*/*mid1* ddPCR assays, droplets with and without *ftsZ* and *mid1* can be distinguished through FAM and HEX amplitude measurements. Droplets are color-coded by target presence: gray (*ftsZ*-/*mid1*-), blue (*ftsZ*+/*mid1*-), green (*ftsZ*-/*mid1*+), and orange (*ftsZ*+/*mid1*+). The number of **(D)** *mid1*-positive and **(E)** *ftsZ*-positive droplets declines across a dilution series in a *ftsZ*/*mid1* ddPCR assay. **(A,B,D,E)** Each dot represents a droplet measured in ddPCR, with gray and blue/green indicating negative and positive, respectively. Samples are separated by vertical dotted lines along the x-axis. **(F)** *ftsZ* and *mid1* concentration measured by *ftsZ*/*mid1* ddPCR strongly correlates with the dilution series. **(G)** Variation between technical replicates is not significantly correlated with the dilution series. Statistical significance: *P* > 0.05 (ns), *P* ≤ 0.05 (*), *P* ≤ 0.01 (**), *P* ≤ 0.001 (***), *P* ≤ 0.0001 (****). All statistical tests are Pearson’s product-moment correlations. The raw data is available in **Data S3**.

With confirmation that the *mid1*-ddPCR assay works at 60°C across a wide range of target concentrations, we proceeded to evaluate the duplex *ftsZ*/*mid1*-ddPCR assay. As expected, the duplex *ftsZ*/*mid1*-ddPCR assay successfully differentiated between four droplet groups (*ftsZ*-/*mid1*-, *ftsZ*+/*mid1*-, *ftsZ*-/*mid1*+, and *ftsZ*+/*mid1*+) when performed with DNA from *w*Mel-bearing *D. melanogaster* (**Fig 2C**). Minor variations in HEX amplitude, observed when *ftsZ* is present or absent, result in two distinct sets of positive and negative droplets on the HEX channel (**Fig 2D**). We hypothesize that this is caused by bleedthrough of high amplitude FAM signals to the HEX channel. Despite these fluctuations, the differentiation between mid1-positive and mid1-negative droplets remains unaffected (**Fig 2C**). To assess the sensitivity of the duplex assay, we performed *ftsZ*/*mid1*-ddPCR on a 1:10 dilution series. Both *ftsZ*- and *mid1*-positive droplets were detected across four orders of magnitude, from 10^−1^ to 10^−4^ (*ftsZ*: 95% CI [488, 514] to [0.14, 0.87] copies/μL; *mid1*: 95% CI [286, 305] to [0.06, 0.60] copies/μL, respectively; **Fig 2D, E**). Dilution factor was strongly correlated with both *ftsZ* and *mid1* concentration (*ftsZ*: 95% CI [0.997, 1] Pearson’s *r^2^*, *P* = 6.90e-11; *mid1*: 95% CI [0.897, 0.997] Pearson’s *r^2^*, *P* = 2.03e-06; **Fig 2F**), suggesting high sensitivity. Additionally, while the coefficient of variation (CV) for both *ftsZ* and *mid1* is slightly higher at lower concentrations, this difference is not statistically significant (*ftsZ*: 95% CI [0.608, 0.987] Pearson’s *r^2^*, *P* = 0.28; *mid1*: 95% CI [0.109, 0.997] Pearson’s *r^2^*, *P* = 0.08; **Fig 2G**), suggesting consistent precision across concentrations. To determine the LoD, we analyzed seven NTCs compared to diluted targets. A single *ftsZ*-positive droplet was detected, and no *mid1*-positive droplets were observed among 124,773 total NTC droplets. Based on the NTC with the widest 95% CI for *ftsZ* (95% CI [0.003, 0.31] copies/μL) and *mid1* (95% CI [0, 0.24] copies/μL), we estimate LoDs of approximately 7 and 5 copies per 20 μL reaction for *ftsZ* and *mid1*, respectively.

### Quantifying a DNA spike-in control with ddPCR

While *Wolbachia* density is a widely used metric in *Wolbachia*-host interaction studies, it has limitations. Density assays often assume a constant number of host genome copies per cell, which may not hold true under varying conditions (Christensen et al., 2019). In such cases, measuring absolute *Wolbachia* abundance, as enabled by the singleplex *ftsZ*-ddPCR assay, may be more informative. However, density assays offer an advantage by normalizing *Wolbachia* DNA against host DNA, mitigating technical variations in DNA purification efficiency by assuming equal recovery of both targets. To improve upon absolute *ftsZ* abundance assays, we developed a duplex *ftsZ*/spike-ddPCR assay that simultaneously quantifies *ftsZ* and a DNA spike-in control added to samples immediately after cell lysis during DNA purification. We employed the DNA Spike I kit from TATAA Biocenter, which includes a synthetic DNA template, primers, and a HEX-labeled probe for qPCR amplification.

For experiments validating spike-ddPCR assays, we added the spike-in control after purification to ensure high target abundance. We evaluated the impact of annealing temperature on the spike-in control oligos as a ddPCR assay, determining that 60°C is suitable for differentiating positive (*x̄* = 4507) and negative (*x̄* = 1138) droplets (**Fig 3A**). The spike-in control assay exhibited a higher incidence of “rain” (droplets between positive and negative clusters) compared to the *ftsZ*- and *mid1*-ddPCR assays. Despite this, our ability to discriminate between positive and negative droplets remained unaffected since there are relatively few droplets between clusters compared to those within clusters. We attribute the increase in rain to the original design of the spike-in control oligos for qPCR, while the *ftsZ* and *mid1* oligos were optimized specifically for ddPCR. We also confirmed that the spike-ddPCR assay is specific to samples containing the synthetic template (**Fig 3B**).

**Figure 3.**
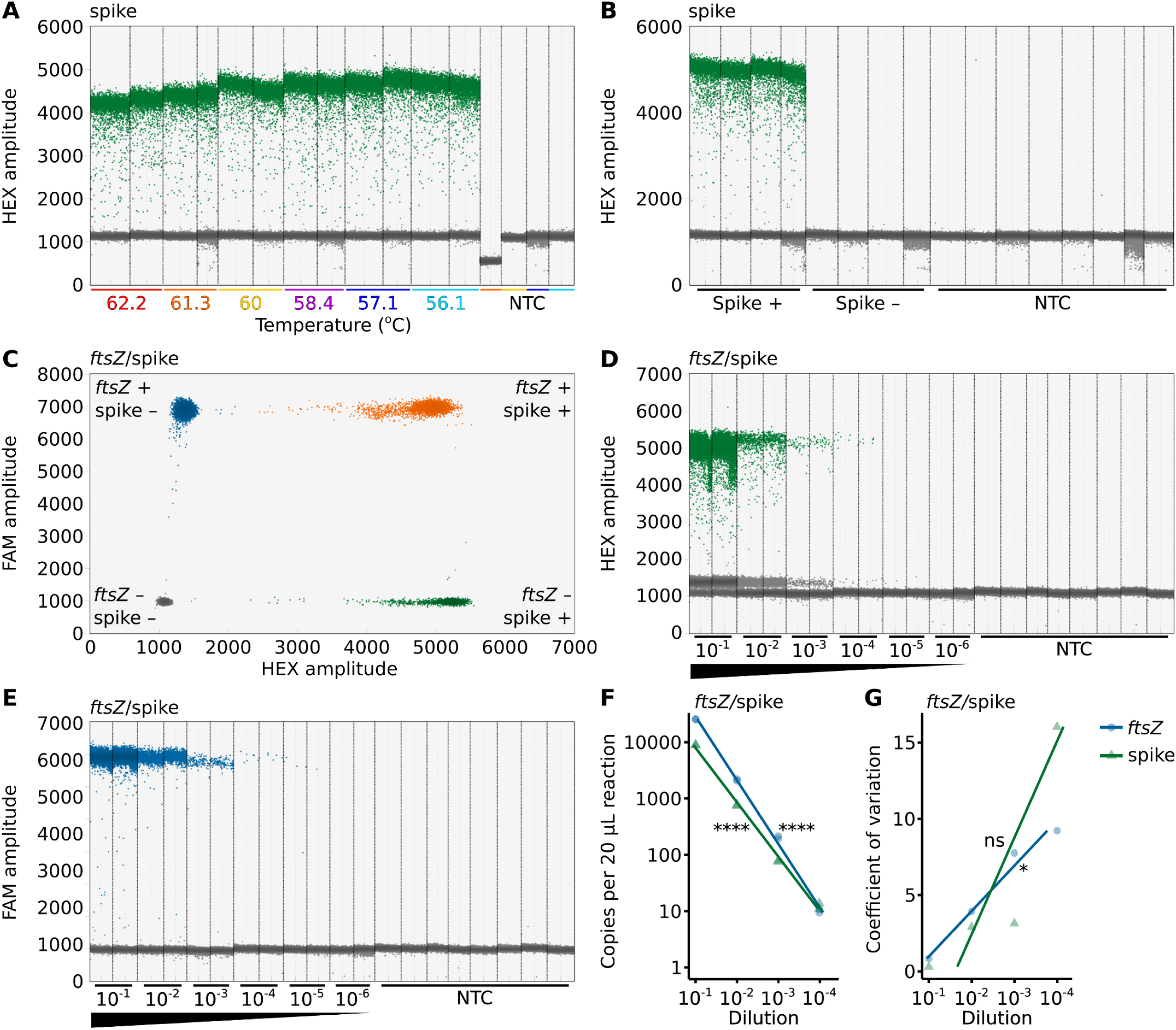
A sensitive *ftsZ*/mid1 ddPCR assay for measuring *Wolbachia* abundance while correcting for DNA purification efficiency. **(A)** Spike detection is consistent across a range of annealing temperatures (62.2°C to 56.1°C). **(B)** Spike detection is specific to samples containing the spike-in control template. **(C)** In duplex *ftsZ*/spike ddPCR assays, droplets with and without *ftsZ* and spike can be distinguished through FAM and HEX amplitude measurements. Droplets are color-coded by target presence: gray (*ftsZ*-/*mid1*-), blue (*ftsZ*+/*mid1*-), green (*ftsZ*-/*mid1*+), and orange (*ftsZ*+/*mid1*+). The number of **(D)** spike-positive and **(E)** *ftsZ*-positive droplets declines across a dilution series in a *ftsZ*/spike ddPCR assay. **(A,B,D,E)** Each dot represents a droplet measured in ddPCR, with gray and blue/green indicating negative and positive, respectively. Samples are separated by vertical dotted lines along the x-axis. **(F)** *ftsZ* and spike concentration measured by *ftsZ*/spike ddPCR strongly correlates with the dilution series. **(G)** Variation between technical replicates is not significantly correlated with the dilution series. Statistical significance: *P* > 0.05 (ns), *P* ≤ 0.05 (*), *P* ≤ 0.01 (**), *P* ≤ 0.001 (***), *P* ≤ 0.0001 (****). All statistical tests are Pearson’s product-moment correlations. The raw data is available in **Data S3**.

Next, we performed duplex *ftsZ*/spike-ddPCR assays, revealing them to successfully differentiate four droplet groups (*ftsZ*-/spike-, *ftsZ*+/spike-, *ftsZ*-/spike+, and *ftsZ*+/spike+) in *w*Mel-bearing *D. melanogaster* samples where the spike-in template was added (**Fig 3C**). As observed in the *ftsZ*/*mid1*-ddPCR assay, *ftsZ* presence in the *ftsZ*/spike-ddPCR droplets leads to minor fluctuations in droplet HEX amplitude (**Fig 3D**). However, these variations do not compromise the ability to distinguish between droplets with and without the spike-in template (**Fig 3C**). To assess the sensitivity of the duplex *ftsZ*/spike-ddPCR assay, we performed a 1:10 dilution series of *w*Mel-bearing *D. melanogaster* DNA spiked with known amounts of the synthetic DNA template. Both *ftsZ*- and spike-in-positive droplets were reliably detected across a range of concentrations (*ftsZ*: 95% CI [1251, 1303] to [0.2, 0.99] copies/μL; spike: 95% CI [432, 458] to [0.28, 1.15] copies/μL; **Fig 3D, E**). Though, we noted that spike abundance is overestimated at a dilution factor of 10^−4^. A strong correlation between the expected and observed concentrations of both *ftsZ* and the spike-in control (*ftsZ*: 95% CI [0.982, 0.999] Pearson’s *r^2^*, *P* = 1.03e-08 ; spike: 95% CI [0.973, 0.999] Pearson’s *r^2^*, *P* = 3.23e-08 ; **Fig 3F**) indicates high sensitivity and accuracy. Additionally, the CV for *ftsZ* is significantly higher at lower concentrations (95% CI [0.243, 1] Pearson’s *r^2^*, *P* = 0.013), while the spike-in control exhibits a marginally higher CV (95% CI [0.323, 0.994] Pearson’s *r^2^*, *P* = 0.135; Fig 3G). Zero false-positive droplets were observed among 264,312 total droplets. Based on the NTC with the widest 95% CI for *ftsZ* (95% CI [0, 0.269] copies/μL) and the spike-in (95% CI [0, 0.269] copies/μL), we estimate LoDs of approximately 6 copies per 20 μL reaction for *ftsZ* and the spike-in control.

## Discussion

*Wolbachia* abundance is an important metric in many studies of *Wolbachia*-host interactions, and is traditionally measured using qPCR. While qPCR is often sufficient, it can be difficult to detect rare targets and is prone to accuracy and precision issues when PCR efficiency is lower than 100% or when it varies across reactions. ddPCR is more sensitive than qPCR and resistant to PCR efficiency variation (Hoshino and Inagaki, 2012; Whale et al., 2012), offering an alternative to quantifying *Wolbachia* gene copies (Fisher et al., 2019; Hickin et al., 2022; Kakumanu et al., 2024; Kilpatrick et al., 2024). In this study, we report three ddPCR assays to count *Wolbachia ftsZ*-gene copies: *ftsZ* ddPCR, *ftsZ*/spike ddPCR, and *ftsZ*/*mid1* ddPCR. Designed to target *ftsZ* sequences from 106 supergroup A *Wolbachia* strains, these ddPCR assays are versatile and may be applicable to a broad range of studies to count *Wolbachia* genomes, including those involving *w*Mel *Wolbachia*. *w*Mel is used to slow the spread of dengue and Zika viruses from mosquitoes to humans (e.g. Utarini et al., 2021).

These *ftsZ*-based ddPCR assays can reliably detect as few as 7 to 12 *Wolbachia* gene copies per 20 µL reaction, making them suitable for quantifying rare *Wolbachia*. We also observed similar LoDs for *mid1* and the spike-in control in duplex assays. We recommend excluding samples with 95% CIs extending below the LoD from further analysis. However, it is crucial to interpret these LoDs with caution. LoDs are determined by two factors: the false-positive detection rate and the ability to reliably detect targets at specific concentrations. In most cases, we report target detection at the limit of the false-positive detection rate, making the LoD equivalent to the upper 95% CI for false-positive detection. However, the 95% CI for false-positive detection can fluctuate between experiments due to contamination and the number of droplets measured in NTCs. Given the sensitivity of these assays, it is essential to conduct ddPCR in a clean environment and take meticulous precautions to prevent sample cross-contamination. As laboratory cleanliness and individual researcher technique can impact false-positive rates, we recommend reporting experiment-specific LoDs based on concurrent NTC reactions.

In addition to *ftsZ*, *mid1*, and spike abundance being reliably detected from rare targets, there was minimal variation between replicate measures across dilutions, indicating that results are reproducible (Devonshire et al., 2023). However, the CV between replicate measures was always higher at lower concentrations. Although this relationship was not statistically significant for most assays, it is possible that the lack of significance stems from insufficient statistical power to detect a correlation between dilution factor and concentration across technical replicates. We interpret these results to suggest that ddPCR precision is marginally lower when quantifying rare targets with these assays. This is expected, as accurate quantification of rare targets is more susceptible to stochastic fluctuations in target distribution within the sample, which can be carried over to the reaction. These fluctuations can lead to minor variations in the number of positive droplets with greater impacts on precision when targets are rare. To improve assay precision for very rare targets, we recommend performing samples in duplicate and treating the results of the duplicate reactions as a single data point (Quan et al., 2018). This approach doubles the target and droplet abundance, increasing confidence in concentration measurements. Despite the variation in precision, the accuracy of the results appears to be unaffected by target concentration, with the exception of the spike-in control, which is slightly overestimated when rare. However, in experimental conditions, rare spike-in targets would indicate a problem in DNA purification efficiency which might warrant exclusion of the sample or reevaluation of processing techniques.

The three ddPCR assays presented in this study have unique advantages and limitations that should be carefully considered before experimental design. The *ftsZ* and *ftsZ*/spike ddPCR assays enable absolute quantification of *Wolbachia* gene copies, providing titer estimates. Titer estimates are useful when the total amount of *Wolbachia* in an insect, tissue, or environment is desired. The *ftsZ* assay is the simplest and least expensive, and is ideal when target abundance in the original sample is not a primary concern, technical variation is evenly distributed across treatment groups, and biological variation between treatment groups is substantial enough to overcome the noise from technical variation. It is advisable to conduct a preliminary experiment to validate these assumptions. If the assumptions are not met, the *ftsZ*/spike ddPCR assay may be a more suitable alternative. The *ftsZ*/spike ddPCR assay requires two key modifications to the standard workflow: a spike-in control is added to samples after cell lysis and before DNA purification, and control reactions are established where the spike-in control is not subjected to extraction conditions but is diluted to the elution volume used for extracted samples. By comparing the spike concentration in samples and controls, it becomes possible to calculate recovery efficiency and adjust *ftsZ* concentrations appropriately.

Titer-based *Wolbachia* abundance assays are useful when the overall quantity of *Wolbachia* in a sample is the primary interest. However, they have limitations, particularly when the number of *Wolbachia* cells per host cell is more relevant than the total *Wolbachia* count. If the number of host cells is consistent across treatment groups (Christensen et al., 2019), titer assays may serve as an approximation of density. However, this assumption can be challenging to validate, and variations in host cell numbers between treatment groups or samples can introduce experimental noise that can obscure important biological signals. The *ftsZ*/*mid1*-ddPCR assay can be useful in such scenarios to normalize *Wolbachia* abundance to the concentration of host genome copies. Notably, this normalization also controls for extraction efficiency variation across samples by assuming that host genome copy number is invariable across samples and that only *Wolbachia* genome copies change. This assumption can be evaluated by comparing *mid1* abundance across treatments before using them for normalization. However, the *ftsZ*/*mid1*-ddPCR assay has several limitations: *mid1* oligos are designed only for *Drosophila*, *Scaptomyza*, and *Zaprionus* species, limiting its utility in other hosts; PCR-based density assays assume a constant host genome copy number per cell, which may not hold true across all experimental conditions, such as dietary variations (Christensen et al., 2019); and while normalizing against *mid1* can account for cross-sample extraction variation, it does not account for variation in DNA purification efficiency which inhibits efforts to determine *Wolbachia* abundance in the original sample. These limitations can be addressed by replacing *mid1* oligos with newly designed oligos targeting the host species of interest, using the *ftsZ*/spike-ddPCR assay if appropriate, or applying microscopy-based assays to directly count *Wolbachia* within host cells.

## Materials & Methods

### Insect lines, care, and maintenance

Experiments were performed using *D. melanogaster* from the *y^1^w\**stock (BDSC 1495), both with and without *w*Mel *Wolbachia*. The *Wolbachia*-free line was generated through three generations of tetracycline treatment in a previous study (Detailed protocol in Cooper and Shropshire, 2024a; LePage et al., 2017) and has been maintained for over seven years since treatment (Ballard and Melvin, 2007). Flies were maintained under a 12:12 light:dark cycle at 23°C within a *Drosophila* incubator (Percival DR-36VL) using standard narrow *Drosophila* vials (Flystuff 32-113RL) containing 7-10 mL of fly food (Detailed protocol in Wheeler et al., 2024). Adult flies were anesthetized with CO_2_ during experiments. *Wolbachia* cytotypes were periodically tested by extracting DNA from pools of three randomly sampled flies from each stock using a squish-buffer method, followed by PCR amplification of the *Wolbachia* surface protein (*wsp*) and the host 28s rDNA (Detailed protocol in Cooper and Shropshire, 2024b).

### Sample collection and DNA extraction

Samples destined for ddPCR assays consisted of randomly selected flies transferred individually to 1.5 mL centrifuge tubes (Eppendorf, 05414203) containing three 2.8 mm ceramic homogenizing beads (VWR, 10158-554). Flies were homogenized immediately after collection using a bead mill (Benchmark Scientific, BeadBlaster 96). DNA was extracted using the QIAwave DNA Blood & Tissue Kit (Qiagen, 69554) (Detailed protocol in Hartman and Shropshire, 2024). DNA sample quality was determined using a NanoDrop One^C^ spectrophotometer (ThermoScientific, 840274200). Samples with A260/A280 or A260/A230 below 1.8 were excluded from further analysis. For annealing temperature and spike-specificity assays, we added 2 µL of a 0.005 ng/µL DNA spike-in control to 100 µL of DNA (TATAA Biocenter, DS25SI). For *ftsZ*/spike sensitivity assays, we added 20 µL of a 0.005 ng/µL DNA spike-in control to 30 µL of DNA. We used DNA lo-bind tubes (Eppendorf, 0030108418) when creating 1:10 dilution series for sensitivity assays.

### Oligo design for ddPCR

For this study, we developed two ddPCR oligo sets, each consisting of two primers and a fluorescently labeled probe (**Table 1**). The first oligo set is FAM-labeled and targets *ftsZ*, a single-copy *Wolbachia* gene that is highly conserved across the genus (Shropshire et al., 2022). The second oligo set is HEX-labeled and targets *mid1*, a single-copy “universally conserved element” in *Drosophila* and its relatives (Kern et al., 2015). To design oligos, we started by extracting the target sequences from NCBI. For *ftsZ*, we extracted genes annotated as *ftsZ* from all *Wolbachia* genomes available through NCBI on November 8, 2023. For *mid1*, we used BLAST (e-value 1e-10) with the *D. melanogaster* sequence as a query on October 23, 2023. Subsequently, we created a multiple sequence alignment (MSA) using MAFFT (v7) with default parameters (Katoh and Standley, 2013). The consensus sequence from the MSA, with variable sites represented as Ns, was then passed through Primer3web (v4.1.0) to generate the oligos (Koressaar et al., 2018). In both cases, the sequence list was iteratively reduced and MSAs were regenerated until oligos meeting our criteria, defined below, could be created. The final aligned sequence lists are available in **Data S1** (for *ftsZ*) and **Data S2** (for *mid1*). The accession numbers for the relevant genomes are listed in **Table S1** (for *ftsZ*) and **Table S2** (for *mid1*). The primers and probe sequences for the DNA spike-in control are proprietary (TATAA Biocenter, DS10PHEXI).

**Table 1.**
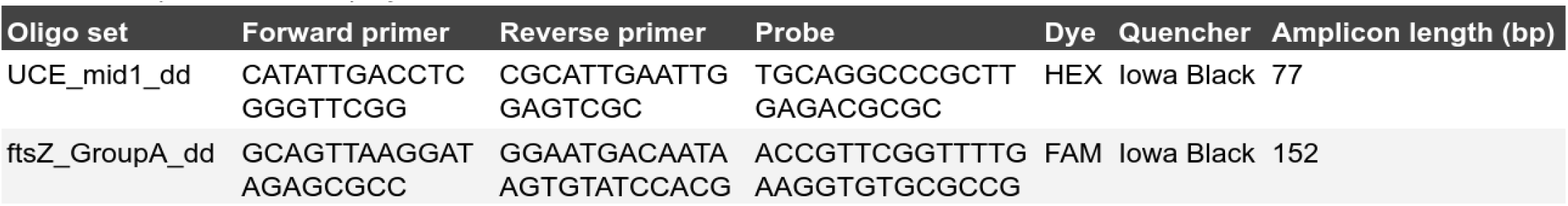
Primers and probes used in this study. The UCE_mid1_dd and ftsZ_groupA_dd oligo sets were used for ddPCR. Sequences are displayed in the 5’ to 3’ orientation.

Our selection criteria for primers included an oligo size of 18–22 bp, a product size between 70–150 bp, a GC content of 40–60%, a GC clamp of 2, a Tm of 58–62°C, a maximum Tm difference of 2°C, and a maximum poly X of 3. Probe selection criteria included an oligo size of 18–30 bp, a Tm of 65–70°C, a GC content of 30–80%, and a maximum poly X of 5. Tm was calculated according to SantaLucia with a monovalent cation concentration of 50 mM, a divalent cation concentration of 3.8 mM, and a dNTP concentration of 0.8 mM (SantaLucia, 1998). Oligos were subsequently tested for specificity using Primer Blast against the human (taxid: 9606), *Drosophila melanogaster* (taxid: 7227), and *Wolbachia* (taxid: 953) genomes. Oligos were purchased premixed from Bio-Rad at a primer:probe ratio of 900 nM:250 nM. Oligos were not diluted before use in ddPCR assays.

### Wolbachia phylogeny

*Wolbachia* genomes used in this study are listed in **Table S1**. To create a *Wolbachia* phylogeny from these genomes, we first identified single-copy orthologs to known bacterial genes using Prokka (v1.14.5) (Seemann, 2014). We excluded pseudogenes, paralogs, and genes with frameshifts or gaps by selecting for genes that were full-length across the genomes of interest. These single-copy full-length orthologs were concatenated and aligned with MAFFT (v7) (Katoh and Standley, 2013). A phylogeny was built from the alignment using RevBayes (v1.1.1) (Hohna et al., 2016), as previously described (Shropshire et al., 2024, 2022).

### ddPCR

All ddPCR reactions were conducted in a 20 µL volume. Singleplex *ftsZ* and *mid1* reactions consisted of 10 µL of ddPCR Supermix for Probes (no dUTP) (Bio-Rad, 1863023), 0.5 µL of the relevant oligo set, 7.5 µL of nuclease-free water, and 2 µL of DNA template. Duplex *ftsZ*/*mid1* reactions were prepared similarly, replacing 0.5 µL of water with 0.5 µL of the second oligo set. Singleplex and duplex DNA spike-in reactions contained 1 µL of a supermix containing forward and reverse primers for the DNA spike-in control and 0.5 µL of a spike-targeting probe. PCR plates were sealed with an adhesive film (Bio-Rad, MSB1001), vortexed for 10 seconds to mix (Four E’s Scientific, MI0101002), and centrifuged for 2 minutes at 2204 x g (Eppendorf, 5430-R). After removing the adhesive film, 19.5 µL of each reaction mixture was transferred to a droplet generation cartridge (Bio-Rad, 1864007), and 70 µL of droplet generation oil for probes (Bio-Rad, 1863005) was added to the adjacent well. A gasket was added (Bio-Rad, 1864007), and the cartridge was placed in a droplet generator (Bio-Rad, QX200) to create droplets. The 40 µL droplet well was transferred to a 96-well plate, sealed with a heat-activated adhesive film (Bio-Rad, 1814040 and PX1), and placed in a thermal cycler (Bio-Rad, C1000 or S1000) for PCR. Reaction conditions varied across experiments, as described in the results section. After PCR, samples were maintained at 12°C until analysis on a droplet reader (Bio-Rad, QX200). Samples were excluded if they yielded fewer than 10,000 droplets. We present a detailed ddPCR protocol for *Wolbachia* abundance assessment on protocols.io (https://www.protocols.io/private/65EC1DB4B26811EFAD000A58A9FEAC02).

### Statistical analysis and figure generation

The QX Manager software (v2.1.0; Bio-Rad) was used to calculate ddPCR target concentration and confidence intervals for and to produce ddPCR plots. Pearson correlation analyses were performed in R (v4.4.1) using RStudio (v2024.04.2). The ‘ggplot2’ package (v3.4.4) in R was used to create correlation plots (R Core Team, 2024; Wickham, 2016). Inkscape (v1.3.2; Inkscape Developers) was used to modify figure aesthetics.

### Data availability

All data are publicly available in the supplement of this manuscript.

## Acknowledgments

We thank Hunter Hill and Tim Wheeler for helpful comments on an earlier version of this manuscript. We also thank Sarah Daniel for ddPCR troubleshooting assistance and Helene Hartman for assistance in the laboratory. Comments by Lore Van Vlaenderen, Helene Hartman, and other members of the Shropshire Lab greatly improved the quality of this work. Support from the staff in Lehigh University’s Department of Biological Sciences is gratefully acknowledged.

## Contributions (CRediT Classification)

Conceptualization: JDS. Data curation: AKN, WRC, JDS. Formal analysis: AKN, WRC, JDS. Funding acquisition: AKN, JDS. Investigation: AKN, FL. Methodology: AKN, FL, JDS. Project administration: JDS. Resources: JDS. Supervision: JDS. Visualization: AKN, JDS. Writing – Original draft: JDS. Writing – Review & editing: AKN, FL, WRC, JDS.

